# Time Regained: How the Human Brain Constructs Memory for Time

**DOI:** 10.1101/186601

**Authors:** Brendan I. Cohn-Sheehy, Charan Ranganath

**Affiliations:** Center for Neuroscience, University of California, Davis, CA, 95618, USA; Department of Psychology, University of California, Davis, CA, 95616, USA; Neuroscience Graduate Group, University of California, Davis, CA, 95618, USA; Physician Scientist Training Program, University of California, Davis, CA, 95817, USA

## Abstract

Life’s episodes unfold against a context that changes with time. Recent neuroimaging studies have revealed significant findings about how specific areas of the human brain may support the representation of temporal information in memory. A consistent theme in these studies is that the hippocampus appears to play a central role in representing temporal context, as operationalized in neuroimaging studies of arbitrary lists of items, sequences of items, or meaningful, lifelike events. Additionally, activity in a posterior medial cortical network may reflect the representation of generalized temporal information for meaningful events. The hippocampus, posterior medial network, and other regions—particularly in prefrontal cortex—appear to play complementary roles in memory for temporal context.

**Highlights:**

- The hippocampus encodes information about temporal contiguity, order, and event structure.
- Posterior medial cortical areas represent order across meaningfully coherent events.
- Prefrontal and subcortical contributions to temporal memory deserve further study.

## Introduction

Human experience unfolds over time, and human memory depends on mechanisms that allow time to be represented and subjectively traversed when replaying past and predicting future experiences [1–3]. Human functional magnetic resonance imaging (fMRI) studies have begun to uncover the neural manifestations of temporal memory [4,5], in parallel with a rich literature involving electrophysiology [6,7] and non-human animals [5,8,9]. Much of this work has implicated the hippocampus, a region known to contribute to episodic memory, in memory for temporal information [4,5,10]. In this paper, we will review recent progress in our understanding of how human memory for temporal information is supported by the hippocampus and by neocortical areas that interact with the hippocampus.

## Temporal Organization in Episodic Memory

Behavioral research has indicated that memories are generally not explicitly “time stamped,” such that people become immediately aware of the specific time at which a recollected event occurred. Instead, considerable evidence suggests that temporal information in episodic memory is reconstructed from retrieved information about items, the environment, and one’s internal state during the event [11]. Friedman [11] added that such reconstructive inferences are easier to perform when one can draw from general knowledge about similar events that have a characteristic temporal or sequential structure. For instance, if asked what time your favorite song was played at a jazz show, you could infer the approximate time through a combination of prior knowledge (e.g., “jazz shows usually include a half-dozen songs, and start around 10pm”) and event-specific information (e.g., “it was mid-show”).

Although episodic memories are not necessarily time-stamped, there is reason to believe that they are *temporally organized*. Specifically, at some level of representation, episodic memories carry information that is specific to a place and time [3], such that events that occurred in close succession are represented in a more similar manner than are events that occurred far apart in time [12–15]. Consistent with this idea, many studies have shown that when one recalls a studied item, there is a higher likelihood of subsequently recalling other items that were in close temporal proximity [16]. By one classic view, this *temporal contiguity* effect might reflect the fact that factors in one’s environment and internal state change gradually over time [17,18]—these factors can be collectively considered as the “temporal context,” and episodic memories may be represented by associating information about specific items to a gradually-changing context representation [12] (i.e., “binding item and context information”). Contextual change may be entirely random [17–19], or it may additionally incorporate information about recently processed items, as suggested by the Temporal Context Model (TCM) [12,14,16]. According to these models, events that occur close together in time are linked through overlap in the lingering neural representation of temporal context [16,20]. Consequently, a recalled event can serve as a cue to retrieve other events from the same time period. As we will describe below, this idea captures many aspects of the temporal organization of representations in the hippocampus and other brain regions.

Although many models operationalize temporal context as a continuous variable, available evidence indicates that this is not the case. As described in more detail in Box 1, people tend to break up a continuous sensory stream into chunks of time that correspond to “events”. In a typical memory experiment, an entire list of items may correspond to a coherent event, and these situations are well-described by temporal context models. In other cases, however, unpredicted changes in the structure of a list, such as changes in a stimulus category[21,22] or encoding task[14] can elicit the perception of boundaries between segments of the list. Other studies have more directly assessed the temporal structure of event memory by studying memories for discrete sequences of events or memories for meaningful, complex events that extend over long timescales. In these cases, knowledge about the temporal structure of similar events can fundamentally shape the neural representation of the event.

#### Box 1: How Events are Constructed and Understood

Bartlett[65] proposed that memories for the past are reconstructed by using structured knowledge about the world called *schemas*, and contemporary theories propose that people use schemas about classes of events (“*event schemas*”) in order to comprehend incoming sensory information [40,41,66–68]. For instance, according to Event Segmentation Theory (EST) [40,41], knowledge from event schemas is retrieved in order to construct an online *event model*, a temporal framework that is used to generate predictions about what will occur next. When the event model is no longer predictive, the current event model is updated, thereby establishing an *event boundary* between the previous event representation and the current event. Event boundaries are reliably triggered by temporal shifts (e.g. in written narratives)[66,69,70] or physical boundaries in a spatial environment [71,72], but they can also be triggered by non-spatiotemporal features like characters, objects, goals, salience, and causality, and their interrelations [41,67]. Critically, event boundaries appear to disrupt the temporal organization of memory. For instance, although retrieval of an item can facilitate retrieval of temporally-adjacent items [14], retrieving items can actually impair retrieval of across-boundary items [66,70]. Furthermore, people are impaired at retrieving the relative order of two items if they were separated by an event boundary [22,27].

In the following sections, we will review results from neuroimaging studies of memory for temporal information and the underlying neural representation of temporal information in memory. Given the influences of event structure on both subjective perception of time and on neural representations of temporal memory, we will separately examine paradigms at different points along the continuum from undifferentiated lists to meaningful, complex events.

## Memories within a Contiguous Timeline

As noted above, the temporal contiguity effect has been used to support models which suggest that episodic memory is organized by temporal context. Given the role of the hippocampus in episodic memory, Kragel et al. [23] tested whether contiguity in the temporal order of recall would be supported by activity in the hippocampus and two closely related medial temporal lobe regions, the parahippocampal cortex (PHC) and perirhinal cortex (PRC). Participants studied lists of words, and they were subsequently scanned during free recall of the studied words. Neural activity during recalled items was used to scale one of two parameters in TCM[14]: temporal reinstatement, or the degree to which a recalled item reinstates its previous temporal context; and retrieval success, or the probability of recalling an additional item. Kragel et al. found that activity in the PHC and posterior hippocampus scaled with temporal reinstatement, whereas activity in the PRC and anterior hippocampus scaled with retrieval success. These results are consistent with the idea that the hippocampus and PHC process information that is analogous to a contiguous representation of temporal context, whereas the PRC may process information about recalled items.

If the PRC supports item representations, and the hippocampus binds item and context information, then these regions could support memory for temporal order in different ways. More specifically, Jenkins and Ranganath [24] proposed that the hippocampus should directly support retrieval of temporal context, whereas the PRC should indirectly support temporal memory if participants use the strength of item representations as a heuristic for recency. To test this prediction, they scanned participants as they performed semantic judgments on an extended continuous sequence of objects, and after scanning, participants judged the relative order of object pairs from within the sequence.

Jenkins and Ranganath hypothesized that the spatial pattern of activity across the hippocampus (i.e., “voxel patterns”)[25] would carry information about temporal context. If so, then changes in hippocampal voxel patterns over time should enable successively presented items to be more distinctive from one another, thereby supporting accurate temporal order memory. Consistent with this idea, larger changes in hippocampal voxel patterns predicted successful temporal order retrieval, and a similar effect was seen in the medial prefrontal cortex, but not in the PRC. Jenkins and Ranganath reasoned that if the strength of an item’s memory representation contributes to assessments of temporal context, then one would predict that PRC activity during encoding should be predictive of perceived item recency. Consistent with this prediction, PRC activation during item encoding was positively correlated with perceived recency at retrieval, and a similar effect was seen in the lateral prefrontal cortex.

These findings suggest that hippocampal activity may represent temporal order in terms of a temporal context, and that PRC activity may underlie temporal attributions for particular items.

## Discontinuities Reconfigure Temporal Organization

Whereas the studies described above relied on tasks that continuously unfolded over a metric timeline, discontinuities in experience may alter the neural representation of temporal information, specifically, unpredicted changes in the continuity of incoming information may affect hippocampal activity and memory for temporal information. For instance, Ezzyat and Davachi [26] scanned participants as they processed objects that were paired with scene contexts that switched over time. Multiple objects were successively paired with the same scene, so that participants would be inclined to treat stimuli that were associated with the same scene as part of the same event. Later, participants were shown pairs of objects and asked to determine whether the objects were studied relatively close or far in time from one another. Consistent with the results of Jenkins and Ranganath[24], Ezzyat and Davachi found that hippocampal activity patterns differed more across objects that were later judged to be far apart in time, consistent with the idea that changes in hippocampal activity patterns over time are related to temporal context representation. Interestingly, however, this effect was specific to pairs of objects that were associated with different scene contexts. For objects that were both paired with the same scene context, activity patterns in lateral occipital cortex were related to subjective measures of proximity in time. This dissociation is surprising, potentially indicating that while changes in the sensory stream over time can support a temporal code in the brain, overall hippocampal activity may sometimes be insensitive to such changes if other contextual variables are constant.

Whereas Jenkins and Ranganath [24] and Ezzyat and Davachi [26] found a relationship between hippocampal pattern *dissimilarity* and the ability to differentiate events in time, DuBrow and Davachi [27] found that hippocampal pattern *similarity*[28] was related to better mnemonic differentiation in time. DuBrow and Davachi [27,28] noted that, in their study, participants were prompted to actively associate successive objects as a sequence, and in this case, similarity in the hippocampal representation across successive items would lead to their linkage within an event context. If this is the case, further results from DuBrow and Davachi [29] indicate that this contextual linkage may involve interactions between the hippocampus and prefrontal cortex. Their results showed that, when objects were linked within the same event context, successful retrieval of items in the correct temporal order was associated with higher functional connectivity between the bilateral hippocampus and the medial prefrontal cortex during encoding.

It is worth considering that boundaries between contexts provide a salient cue to signal the passage of time. The findings of Ezzyat and Davachi[26] and DuBrow and Davachi[29] show that hippocampal activity is sensitive to event boundaries, and that these responses are associated with successful subsequent retrieval of temporal information. Nonetheless, activity within smaller areas of the hippocampus may still differentiate between more specific moments in time within a contiguous context [24]. As we will discuss further, this suggests that the hippocampus supports the representation of time in a manner that corresponds to the temporal structure of each particular situation.

## Events as Structured Temporal Sequences

As noted above, real events tend to have a predictable structure—for instance, in an American wedding, typically the bride and groom walk down the aisle before they begin their vows. A minimal way to study memory for temporally structured events is to examine processing of items in a deterministic or probabilistic sequence. Many fMRI studies have shown that hippocampal activity differentiates between learned and unlearned sequences, even when item and spatial information is matched[30–33]. Hippocampal voxel activity patterns, in turn, carry information about the temporal order of items in a sequence [31,34,35]. Hsieh et al. [34] scanned participants while they made semantic decisions on a continuous stream of objects that were either in fixed sequences or in a randomized sequence. Although there were no demands for explicit retrieval during scanning, participants made faster semantic decisions about objects in fixed sequences than for objects in random sequences. Moreover, responses were considerably lagged for the first object in each sequence, suggesting that participants segmented (see **Box 1**) the continuous stream of objects into discrete five-object sets (**Figure 1A)**.

**Figure 1:**
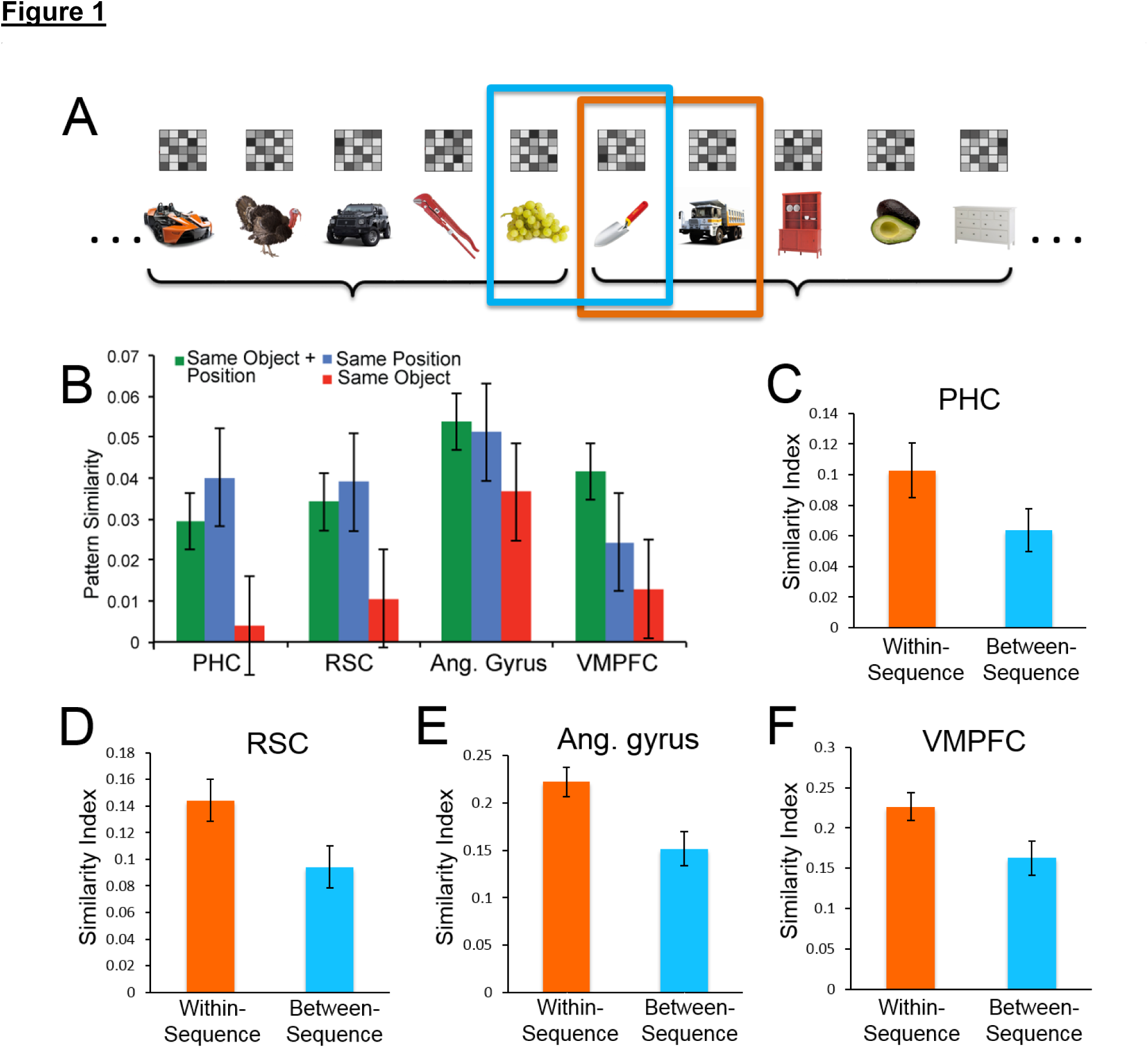
Posterior medial network activity patterns represent structured temporal sequences. **(A)** Experimental task from Hsieh and Ranganath[35]: Participants were scanned while processing a continuous stream of five-object sequences. Boxed items are “within-sequence” (orange box) and “between-sequence” (blue box), respectively. **(B)** Voxel pattern similarity for repetitions of the same object in learned sequence (green bars) and for trials that shared either the same position (blue) or same object in random sequences (red). Values are shown for regions within the posterior medial (PM) network [37], including parahippocampal cortex (PHC), retrosplenial cortex (RSC), angular gyrus (Ang. gyrus), and ventromedial prefrontal cortex (VMPFC). **(C-F)** PM network activity patterns at boundaries between sequences (Frank Hsieh and Charan Ranganath, unpublished data): Activity patterns were more similar between adjacent items in the same sequence (“within-sequence” pairs, orange bars) than between adjacent items in different sequences (“between sequence” pairs, blue bars). **(C**) Parahippocampal cortex. **(D)** Retrosplenial cortex. **(E)** Angular gyrus. **(F)** Ventromedial prefrontal cortex.

The hippocampus showed highly similar patterns of activity across repetitions of objects in the learned, fixed sequences, but not across repetitions of the same objects in random sequences[34]. Hsieh and Ranganath[35] also examined neocortical activity within a previously-characterized, posterior medial (PM) network that is known to support memory-guided behavior through interactions with the hippocampus [36–39]. PM activity appeared to represent information about the temporal position of an object, but these regions generalized across different objects in different sequences[35] (**Figure 1B**). Therefore, both the PM network and the hippocampus carried information about the temporal structure in sequences, but only the hippocampus was additionally sensitive to the identity of specific objects in learned sequence positions. Furthermore, both the hippocampus and regions in the PM network (**Figure 1C-F**) exhibited sharp transitions in activity patterns between sequences, suggesting that PM network regions were sensitive to boundaries between sequences. The results from Hsieh et al. [34] and Hsieh & Ranganath[35] indicate that the PM network may encode general information about structured events, in a manner similar to an *event schema*[40,41] (see **Box 1**), whereas the hippocampus may encode event-specific representations of items and their temporal context.

Complementary results have been reported in studies that have investigated probabilistic sequence relationships. For instance, one study[42] showed that repeated sequential presentation of pairs of objects led to increased similarity in hippocampal activity across the paired objects. Following up on these findings, Schapiro et al.[43] scanned participants after they were exposed to streams of objects with a complex temporal structure. The study was designed such that specific groups of objects were clustered into “temporal communities,” such that presentation of one object would accord with a high likelihood of subsequent presentation of another object from the same community, whereas objects from another community would have a lower likelihood of subsequent presentation. Following learning of the community structure, hippocampal activity patterns were more similar for objects from the same temporal community than for objects that were from different communities.

## Temporal Structure in Complex Lifelike Events

Although studies of sequence memory capture the temporal structure of real events, real-life events are different in the sense that people draw upon knowledge about particular classes of events (i.e., event schemas; **Box 1**), to understand and encode real-life events. Whereas studies of arbitrary sequences repeatedly elicit hippocampal activity, studies of meaningful episodes seem to indicate that the PM network represents temporal structure at longer timescales [44,45].

For example, Chen et al. [46] scanned participants while they viewed an hour-long television show, and as they attempted to recall the show from memory. Regions in the PM network, including PHC, angular gyrus, posterior cingulate, precuneus, and retrosplenial cortex exhibited scene-specific activity patterns, meaning that a consistent pattern of activity was evoked throughout each event in the video. Critically, the order of scene-specific activity patterns observed during viewing was recapitulated as people recalled the show **(Figure 2A-B)**, despite the fact that viewing and recall occurred over different temporal intervals **(Figure 2B)**. This finding indicates that the sequence of activity patterns observed in the PM network was not driven by sensory information, but rather by internal representations of the sequence of events depicted in the video. In a re-analysis of this data, Baldassano et al. [47] showed that regions in the PM network showed abrupt shifts in activity patterns at boundaries between scenes, consistent with what might be expected if these regions were involved in maintaining and segmenting events (**Figure 2C**). Moreover, the correlation between PM network activity and subsequent hippocampal activity predicted later recall success[47]. Taken together, these findings illustrate that the PM network represents meaningful information about sequences of long-timescale events.

**Figure 2:**
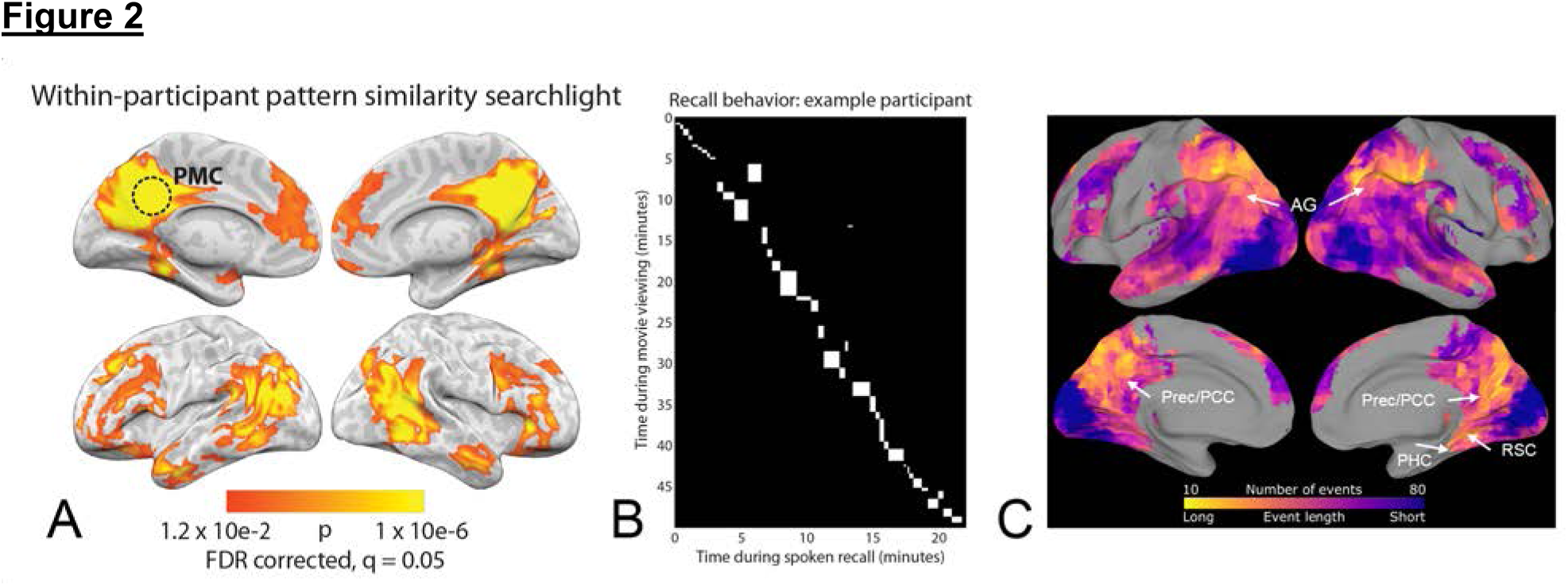
Posterior medial network activity patterns recapitulate naturalistic event sequences. Posterior medial (PM) network activity evoked during a naturalistic paradigm during film viewing and recall, as revealed by searchlight analyses [73]. **(A-B)** Reprinted by permission from Macmillan Publishers Ltd.: *Nature Neuroscience*, Volume 20, Chen, Janice et al., Shared memories reveal shared structure in neural activity across individuals, Pages 115-125, Copyright 2017. **(A)** Scene-specific PM activity patterns are significantly correlated between initial viewing and temporally-ordered recall of the same scenes from an episode of *Sherlock* [46]. PMC = posterior medial cortex. **(B)** Scene durations for viewing vs recall in a representative participant [46]. Each white rectangle represents a particular scene, where width equals the duration of recall and height represents the duration of viewing that scene. **(C)** Posterior medial regions represent events at a coarse timescale [47]. A Hidden Markov Model was used to identify neural event representations at fine and coarse timescales. The colormap, which illustrates the optimal timescale observed for each brain region, shows that regions in the PM network (white arrows) represent events at the longest timescales (warmer colors), on the order of minutes: precuneus/posterior cingulate cortex (Prec/PCC), angular gyrus (AG), parahippocampal cortex (PHC), and retrosplenial cortex (RSC). Reprinted from *Neuron*, Volume 95, Issue 3, Baldassano, Chris et al., Discovering event structure in continuous narrative perception and memory, Pages 709-721.e5, Copyright 2017, with permission from Elsevier.

Further insights have come from studies that have examined how people construct temporally-structured narratives across separate experiences. In these studies, brain activity is examined during processing of a stream of audio or film clips, from which the content of temporally separate clips can be meaningfully integrated into a coherent sequence of events. One such study [48] found that activity in the PM network was higher in magnitude during processing of coherent audio clips when compared with clips that did not comprise a coherent sequence of events (incoherent). Milivojevic et al. [49] compared brain activity patterns as participants watched a continuous movie in which odd-numbered and even-numbered scenes depicted two different coherent narratives involving the same characters and spatial contexts. Activity patterns in the hippocampus and various cortical and subcortical areas carried information about characters and contexts common to the two narratives. However, hippocampal activity patterns elicited by scenes diverged between the two narratives over the course of viewing, as participants acquired more knowledge about the two storylines, despite the fact that these two narratives overlapped in time. This evidence suggests that the hippocampus has the capacity to represent sequences of events within the context of meaningful narratives in a manner that can override a purely metric timeline.

Because the passage of time is often accompanied by changes in spatial location[50], some studies have attempted to disentangle representations of spatial and temporal context. Studies using virtual reality have shown that the hippocampus and PHC may represent interactions between temporal and spatial information [51], and that hippocampal activity patterns in particular correlate with accurate judgments of spatial and temporal distance [51,52] and temporal order [53]. For instance, Deuker et al. [51] scanned participants as they viewed a series of objects that were previously encountered during exploration of a virtual environment. Results showed that hippocampal voxel patterns became more similar for objects that were in close spatial or temporal proximity during the navigation episode, but they became further differentiated for objects that were far apart in space and time. Although the results seem to indicate a central role for medial temporal lobe regions in both spatial and temporal processing, these regions may exhibit different patterns of functional connectivity with frontal and parietal areas depending on whether spatial or temporal information is currently relevant[54].

Nielson et al. [55] observed parallel findings in a study of memory for real-life experiences. GPS-enabled smartphones were used to track participants’ experiences over the course of a month, and then participants were scanned while attempting to recollect events cued by pictures from the smartphones. Hippocampal activity patterns were more similar across pairs of pictures that were taken in relatively close spatial and temporal proximity than across pictures corresponding to events that were far apart in space and time. This evidence suggests that the hippocampus organizes memories for real-life experiences according to their spatiotemporal context, and that this can be seen even across a relatively long timescale.

## Consistent Themes and Questions for Future Research

The evidence reviewed here demonstrates that the hippocampus is clearly involved in memory for temporally specific events[5]. The magnitude of hippocampal activity is correlated with accurate memory for temporal context, and hippocampal voxel patterns carry information about the temporal context of events. These characteristics point to a central mechanism by which the hippocampus can support episodic memory. Specifically, the hippocampus may assign different representations to separate events that occurred at different times, even if the events are otherwise similar. These findings are consistent with the emerging view that the hippocampus could intrinsically organize memories, possibly through neural populations whose activity drifts over time[8,56]. On the other hand, the fact that hippocampal activity is sensitive to event boundaries, ordinal position information, and narrative structure (**Box 1**) suggests that the hippocampus does not encode time in a purely metric sense.

A second theme to emerge is that activity in the PM network[36,37] is elicited in studies of meaningfully structured events. For instance, the PM network appears to process information about narratives or movies across long timescales[44,46–48], PM activity is sensitive to narrative coherence[48], and sequences of PM network activity patterns that are observed during meaningful events are recapitulated when these events are recalled [46,47]. Additionally, the PM network appears to represent ordinal positions within sequences, independent of the items occupying those positions[35]. Given that the PM network interacts closely with the hippocampus, it is possible that these regions represent events in a complementary fashion.

To understand the relative roles of the hippocampus and PM network in temporal memory, it is helpful to consider the broader memory literature. Results from many studies have supported the idea that the hippocampus integrates information about specific items with information about the context in which they were encountered[57–59], and many findings support the idea that the PM network represents knowledge about events, or event schemas[36,37]. Drawing from these models, we propose that, as people process a meaningful event, the PM network may specify the temporal, spatial, and situational relationships between people and things (i.e. an *event model*; see **Box 1**). Accordingly, during experiences that accord with a learned event schema, the PM network activity should provide a framework for temporal relationships amongst elements of an event, and this frame should fundamentally shape event encoding by the hippocampus. Specifically, we would expect the hippocampus to encode a “tag” specifying the currently active event schema, event boundaries, and predictable ordinal or metric temporal relationships within the boundaries of the currently active event. Hippocampal activity patterns should accordingly be more similar between items or events that correspond to similar tags, or more dissimilar when those tags are different, whereas PM network activity should be more similar between events that share similar event schemas. Conversely, when no learned event schema applies to a particular experience, hippocampal activity patterns will encode temporal context in a more continuous manner, reflecting the fluctuation of various stimulus features over time rather than specific positions within a temporal framework.

There is reason to speculate that the medial prefrontal and entorhinal cortex could mediate the relationship between the PM network and hippocampus in representing temporal context in different situations [9,24,29,43,60,61]. The studies reviewed here also highlight roles for other regions in temporal processing, including lateral prefrontal[13,24,29,60–63] and subcortical areas [35,49]. In particular, the lateral prefrontal cortex appears to represent attributions of recency regardless of retrieval success [24], temporal coordinates or order not construed within particular events [13,29], and the presence of mutually predicting temporal regularities among items[43,61]. Further work is needed, however, to determine whether these findings merely reflect a role for prefrontal cortex in cognitive control, or whether they reflect processes that are more specifically relevant for temporal memory.

The time is ripe for human memory. Dynamic, evolving methods continue to advance our network-level understanding of temporal memory, and while physicists continue to debate the nature of time[64], we may be closer to grasping how a material brain can underlie our mental time travel.

## Acknowledgments

We thank Frank Hsieh for sharing unpublished data, Chris Baldassano and Janice Chen for sharing published figures, and Jordan Crivelli-Decker, Halle Dimsdale-Zucker, Marika Inhoff, and Eda Mizrak for useful discussions. This paper was supported by a Vannevar Bush Faculty Fellowship from the Department of Defense (C.R.), a Floyd and Mary Schwall Medical Research Fellowship (B.I.C.S.), and a Neuroscience Graduate Program Fellowship from the University of California, Davis (B.I.C.S.).

